# MARVEL: Microenvironment Annotation by Supervised Graph Contrastive Learning

**DOI:** 10.1101/2024.10.29.620828

**Authors:** Yan Cui, Hongzhi Wen, Robert Yang, Xi Luo, Hui Liu, Yuyin Xie

## Abstract

Recent advancements in *in situ* molecular profiling technologies, including spatial proteomics and transcriptomics, have enabled detailed characterization of the microenvironment at cellular and subcellular levels. While these techniques provide rich information about individual cells’ spatial coordinates and expression profiles, extracting biologically meaningful spatial structures from the data remains a significant challenge. Current methodologies often rely on unsupervised clustering followed by cell type annotation based on differentially expressed genes within each cluster and most of the time will require other information as the reference (e.g., HE-stained images). This is labor-intensive and demands extensive domain knowledge. To address these challenges, we propose a supervised graph contrastive learning framework, MARVEL. MARVEL is a supervised graph contrastive learning method that can effectively embed local microenvironments represented by cell neighbor graphs into a continuous representation space, facilitating various downstream microenvironment annotation scenarios. By leveraging partially annotated examples as strong positives, our approach mitigates the common issues of false positives encountered in conventional graph contrastive learning. Using real-world annotated data, we demonstrate that MARVEL outperforms existing methods in three key microenvironment-related tasks: transductive microenvironment annotation, inductive microenvironment querying, and the identification of novel microenvironments across different slices.

## 1 Introduction

The tissue microenvironment (ME), a dynamic mix of cellular and non-cellular components, forms a complex regulatory network, essential for maintaining the homeostasis of a tissue [2]. The rapid development of *in situ* molecular profiling techniques, such as spatial proteomics (e.g., CODEX [4] and SAFE [12]) and transcriptomics (e.g., Visium [13], Xenium [10], CosMX [7], GeoMX [16], MERFISH [14]), has enabled high-resolution characterization of tissue MEs at both cellular and subcellular levels. These technologies generate detailed spatial coordinates and expression profiles for individual cells, which can be used to annotate cell types and study cell-cell interactions. However, capturing biologically meaningful and disease-relevant spatial structures from those data remains a significant challenge. Existing methods rely on unsupervised clustering combined with differential expression marker analysis, which requires expert interpretation—an approach that is both time-consuming and costly [3, 9, 17].

With the explosive growth of spatial omics data, traditional clustering methods followed by manual annotation becomes increasingly infeasible due to the extreme time and difficulty involved. It is essential to develop models that can capture spatial tissue architectures across slices/samples and automatically assign microenvironment (ME) labels to new slices with minimal prior annotation information. Several previous studies [17, 9] have attempted to tackle cross-slice annotation through unsupervised spatial domain co-clustering to achieve consistent annotations across multiple slices. While these methods address consistency and scalability issues in multi-slice applications, they still suffer from the short-comings of unsupervised clustering: they cannot utilize previous annotations and require manual annotation to translate clustering into biologically meaningful categories.

To bridge the gap between the growing demand for cross-slice annotation and the lack of efficient methods, we propose a new framework, MARVEL (<u>M</u>icroenvironment <u>A</u>nnotation by supervised g<u>R</u>aph contrasti<u>VE</u> Learning), which facilitates label transfer and querying efficiently and flexibly in various scenarios. Inspired by the microenvironment modeling approach from SPACE-GM [18], we leverage cell-neighbor graphs to represent the MEs of each cell. Here, an ME is defined as neighboring cells surrounding a centric cell at various radii, which forms a spatial neighbor graph.

MARVEL is a supervised graph contrastive learning method, i.e., a combination of supervised contrastive learning [11] with graph contrastive learning [21, 20, 5] in a semi-supervised setup. This advanced framework can embed the input cell-neighbor graph into a continuous and representative space while maintaining ME-related information by making full use of limited annotated samples. Meanwhile, MARVEL leverages partially annotated samples as strong positives, improving the quality of representations and mitigating the negative effects of false positives often seen in standard graph contrastive learning. Unlike previous methods focused on extracting aggregated ME embeddings to perform slice-level predictions [3, 9], we propose concentrating on a fine-grained cellular level. The main difference between MARVEL and previous graph-based ME models, e.g., SPACE-GM, lies in their target applications: MARVEL defines each cell’s ME through its cell-neighbor graph and reframes the ME annotation task as a graph classification problem for these cell-neighbor graphs, addressing the cell-level task. Meanwhile, SPACE-GM conducts slice-level predictions, treating ME embedding as an intermediate representation to achieve a final slice embedding.

MARVEL is the first method capable of semi-supervised representation learning for microenvironments in spatial omics data, achieving efficient and precise label transfer for inter- and intra-slice microenvironment annotation with limited label information. Specifically, MARVEL requires only a small amount (0.05 %) of partially annotated ME data from some slices, along with the remaining unannotated slices, as input. It can transfer these limited annotations to both the partially annotated and unannotated slices, demonstrating its transductive label transfer capability (around 80% F1 score). Additionally, thanks to the high quality of learned representations, MARVEL can be used for inductive label annotation and novel ME discovery without retraining, unlike traditional label transfer methods that require co-training on both labeled and unlabeled data. More specifically, in an inductive setting, the microenvironment (ME) to be annotated can be directly assigned a label by querying the pretrained reference representations, while in a traditional transductive setting, the ME to be annotated must be co-trained with the annotated MEs in a semi-supervised manner. Extensive benchmarking tests demonstrate that MARVEL outperforms baseline techniques across three key ME-related tasks: transductive ME annotation, inductive ME query, and novel ME discovery across multiple datasets in all evaluation metrics, indicating the generalization and application potential of MARVEL.

## 2 Method

The input to MARVEL consists of multiple cell-neighbor graphs derived from *N* input spatial omics slices, collectively denoted as 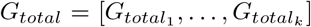, where *k* ∈ { 1, … , *N*} . Each slice contains multiple cells represented as nodes in a graph, with a total of *M* cells in whole slices. Each spatial graph captures the spatial neighborhood relationships between cells in a slice using an adjacency matrix. We denote the spatial neighbor graph by an adjacency matrix *A*_*k*_ ∈ ℝ ^*n×n*^, where *A*_*kij*_ = 1 indicates that cell *i* and cell *j* are adjacent in slice *k*, while *A*_*kij*_ = 0 indicates they are not neighbors.

For each cell *i* within a spatial graph 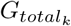, where *i* ∈ { 1, … , *M*}, we first extract a 2-hop cell-neighbor graph as the input to MARVEL. The adjacency matrix of the 2-hop neighbor graph is defined as follows:

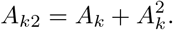

The node features are represented by the matrix *X*_*k*_ ∈ ℝ ^*n×d*^, where *d* is the number of features associated with each cell. Categorical features, such as cell types, can be encoded as a one-hot embedding matrix, while numerical features, such as profiled molecular expressions, are stored as is. The constructed graph representation for each 2-hop neighbor subgraph centered on cell *i* and its index on corresponding slice *k* as *k*(*i*) is expressed as:

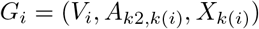

where *V*_*i*_ denotes the set of nodes in the subgraph centered on cell *i* with node number |*V*_*i*_| = *n*_*i*_ , *A*_*k*2,*k*(*i*)_ is the 2-hop adjacency matrix for the subgraph, and *X*_*k*(*i*)_ is the corresponding node feature matrix for the subgraph. The final input dataset for MARVEL consists of the combined graph set {*G*_1_, *G*_2_, … , *G*_*i*_, … , *G*_*M*_}.

The GNN encoder processes the cell-neighbor graphs for input cells. For cell-neighbor graph *G*_*i*_, we input it into the GNN encoder to produce the node embeddings *z*_*j*_. This is formalized as follows:

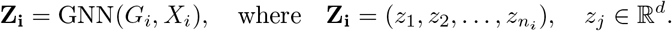

Given the updated node embeddings *z*_*j*_ in *G*_*i*_, we then compute the microenvironment (ME) embedding **h**_*i*_ for the *i*-th cell-neighbor graph by applying mean pooling over the node embeddings. This aggregates the individual node-level representations into a single microenvironment embedding as follows:

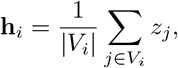

where |*V*_*i*_| denotes the number of nodes in the graph, and **h**_*i*_ ∈ℝ ^*d*^ represents the ME embedding for the *i*-th cell incorporating its neighbor information. The mean pooling operation computes the average of all node embeddings to generate **h**_*i*_, capturing the characteristics of the entire microenvironment.

MARVEL is designed to learn a robust and representative cell-neighbor graph latent representation. Thus, we apply the InfoNCE [1, 6] loss to the ME embedding **h**_*i*_. This loss is designed to maximize the similarity between representations of positive pairs (such as augmented versions of the same ME) while minimizing the similarity between negative pairs, which has shown good performance in previous Graph Contrastive Learning work[21, 20].

The InfoNCE loss is defined as follows:

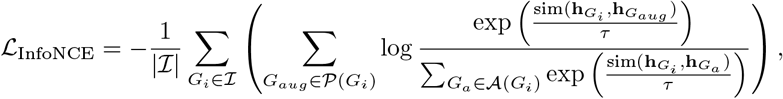

where *ℐ* is the set of all ME graphs in the batch, the temperature parameter *τ* is a scaling factor that controls the sharpness of the similarity distribution, *𝒫* (*G*_*i*_) represents the set of positive ME graph for ME graph *G*_*i*_ (augmented versions), while *𝒜* (*G*_*i*_) denotes the input original ME without data augmentation in the batch, which may include both same-category and differet category samples. The ME representation for *G*_*i*_ is denoted as 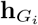, and 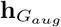 is the representation for the augmented microenvironment graph. The similarity between two ME representations is measured using a similarity function, such as cosine similarity:

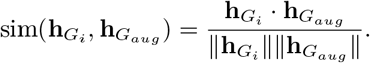

In this InfoNCE loss function, the augmented version *G*_*aug*_ is treated as a positive sample, while the remaining samples within the same batch are treated as negative samples. Furthermore, we apply a supervised contrastive loss[11] to the ME representations **h**_*G*_ for the labeled ME graph. This loss is designed to encourage the ME representations with the same label (referred to as positive pairs) to be more similar, while making the representations of microenvironments graph with different labels (negative pairs) more dissimilar. The supervised contrastive loss is defined as follows:

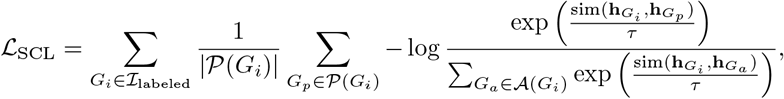

where *ℐ* _labeled_ is the set of all labeled ME graphs in the batch, 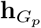 and 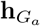 are the representations for positive and input original MEs, respectively.

The final loss function for MARVEL combines both the InfoNCE loss and the supervised contrastive loss:

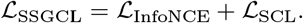

This formulation allows the model to effectively utilize both unlabeled data through augmentations and labeled instances, thereby improving the overall robustness and generalization of the ME graph representations learned.

## 3 Experiment

### 3.1 Dataset preparation, preprocessing and Implementation details

We collected three publicly available large-scale CODEX datasets with pathologist-annotated ground truth MEs to evaluate our proposed method against existing approaches. The first dataset [4] consists of multiplexed immunofluorescence imaging data from 140 tissue regions collected from colorectal cancer (CRC) patients, yielding 140 slices. The second dataset is collected from a TNBC (triple-negative breast cancer) patient and contains 12 slices [15], while the third dataset is from a melanoma patient, comprising 24 slices [8]. For our experiments, we utilize half of the slices from each dataset for training, applicable to both transductive and inductive settings, and reserve the remaining slices for evaluation in the inductive query setting. Specifically, in the transductive learning setting, both labeled and unlabeled instances are trained together; however, label information is only available for the labeled instances, which is often referred to as ‘label transfer.’ In the inductive setting, the annotation of the unlabeled data is directly assigned by the trained model without any co-training procedure.

In all datasets, we stick to the same backbone GNN architecture, the Graph Isomorphism Network (GIN) [19], with identical hyperparameters across experiments. For the transductive label transfer task, we use label proportions of [0.01, 0.05, 0.1, 0.2, 0.3, 0.4] as the seed ME, and evaluate performance using both the cell type and panel expression value-based feature versions of MARVEL. The data augmentation techniques applied during contrastive learning include edge dropout and node feature masking, using a fixed ratio.

In the inductive ME label query and novel ME label identification tasks, we set the default label proportion parameter to 0.1 to construct the reference atlas. Due to the absence of prior methods tailored for these tasks, we use a conventional classifier with the same input features as a baseline. Specifically, we employ cell-type one-hot encoding along with the panel profile values of the central cell, the mean panel expression values, and the cell proportions of neighboring cells as input features for Random Forest and logistic regression models. To ensure a fair comparison, baseline methods and MARVEL use the same set of 2-hop neighboring cells as input.

To ensure the fair benchmarking comparison and remove the variance introduced by model architecture, we apply the same GIN architecture, with supervised classification as the primary training objective. In the inductive ME label query and novel ME label identification tasks, we use reference embeddings based on neighbor cell proportions and panel mean values as the baseline.

### 3.2 Transudtive ME label transfer

Annotating MEs on reference spatial omics datasets often requires significant time and effort from pathologists. As a result, transferring labels from a small subset of annotated cells to unlabeled ones within a slide, also known as transductive label transfer, is a practical and efficient strategy. However, no existing ME annotation method has been designed for this transductive learning task. To tackles this challenge, we propose a graph-supervised contrastive learning framework. We first employ graph-supervised contrastive learning, followed by linear probing to obtain classification results. Specifically, we train a logistic regression model using the labeled embeddings and perform inference on the unseen samples. To evaluate performance, we choose the F1 score (macro F1 score) to address class imbalances and assign equal weights to each cell type in the overall score. The results in Fig 2 demonstrate the superior performance of MARVEL over other benchmark methods.

**Fig. 1:**
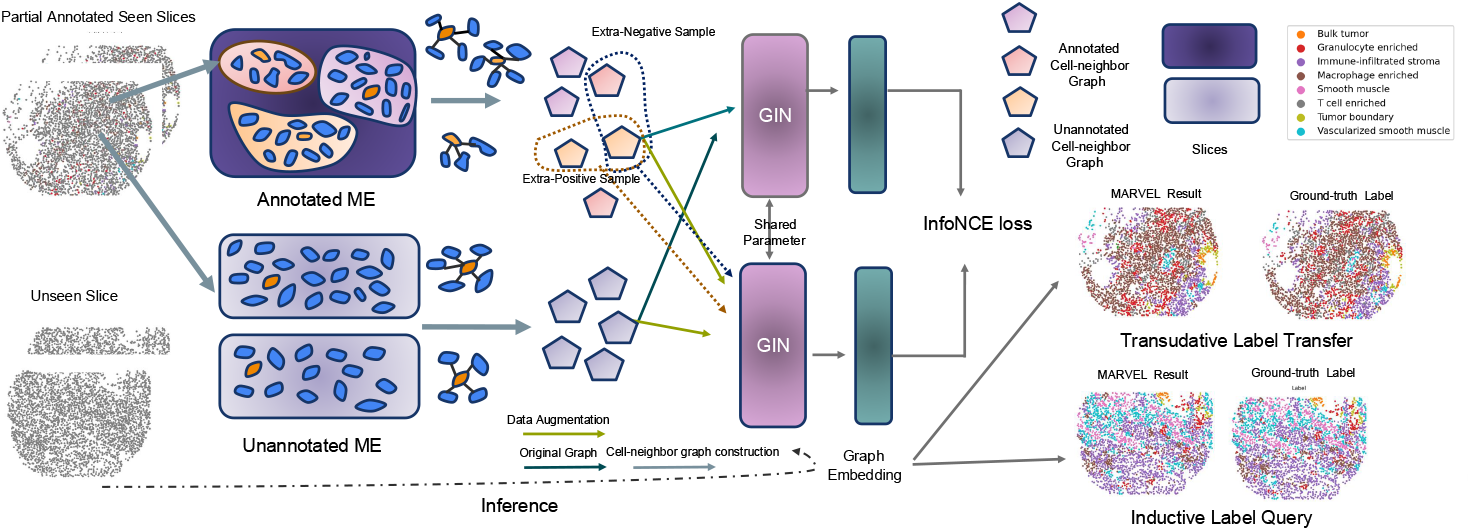
Proposed pipeline for MARVEL: The input to MARVEL includes a small proportion of annotated microenvironments (for each centric cell) and the remaining unlabeled microenvironments from partial annotated seen slices as training slices. These MEs are represented using a 2-hop cell-neighbor graph. The ME representation is constructed through supervised graph contrastive learning with InfoNCE loss. The orange cell and its neighboring cells illustrate an example cell-neighbor graph. Different background colors in the annotated MEs represent corresponding labels. The learned GIN encoder can be utilized to both transducitive label transfer on partial annotated seen slices and inductive label query on the unseen slices.

**Fig. 2:**
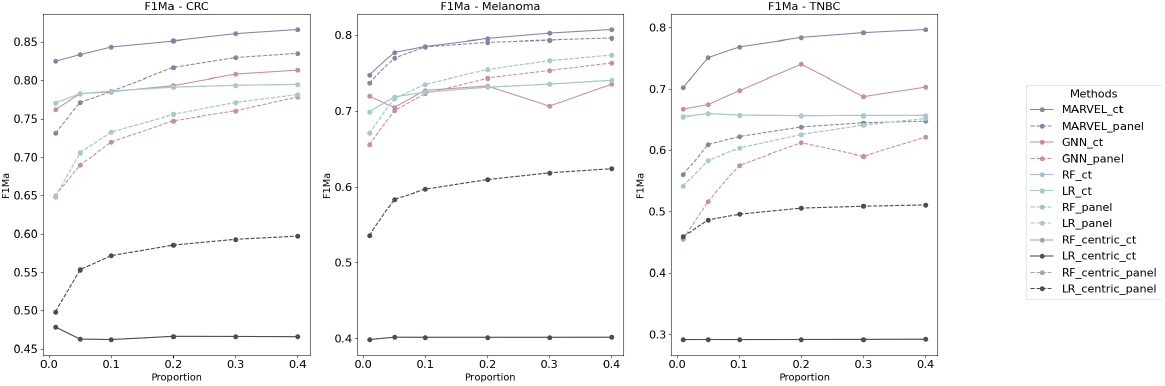
Transductive label transfer results (F1-Macro) across multiple baselines for different label proportions [0.01, 0.05, 0.1, 0.2, 0.3, 0.4] on three different datasets.

**Fig. 3:**
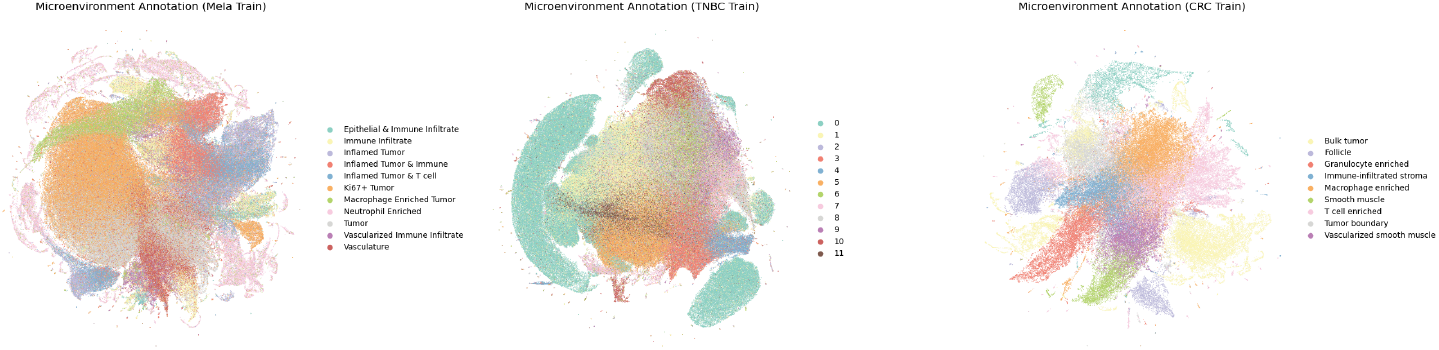
UMAP visualization of the transductive label transfer learned representation with 0.1 labeled proportion.

### 3.3 Inductive ME label query

When dealing with large-scale labeled datasets and an increasing number of new datasets, the transductive method becomes costly. This is because handling a rapidly updated query dataset requires the user to re-train the model each time a new query dataset is introduced. To address this issue, a direct *Inductive Label Query* method, which learns a general classification function on the preconstructed embedding reference dataset, can alleviate the computational cost.

Formally, given a pre-constructed dataset *X*_train_, with a new input query dataset *X*_test_, the label for each sample in *X*_test_ is assigned based on the cosine similarity to the existing embeddings in *X*_train_.

The cosine similarity between a query sample 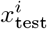 and a reference sample 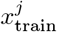 is computed as follows:

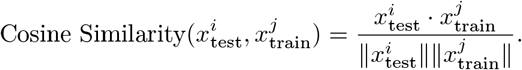

The label for the query 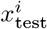 is assigned to the label of the nearest sample in the reference dataset *X*_train_. This inductive method reduces the computational cost by avoiding repeated training by using the pre-constructed reference embeddings.

For evaluation, we calculate the Top-K Hit Ratio, which evaluates whether the true label of a query sample is included among the top K most similar samples in the reference dataset. The Top-K Hit Ratio is defined as:

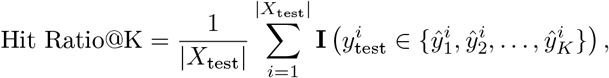

where 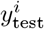 is the true label of the *i*-th test sample, and 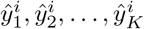 represent the predicted labels of the top K most similar samples from the training dataset. The indicator function **I**(*·*) returns 1 if the true label is within the top K predictions, and 0 otherwise. Higher Hit Ratio@K indicates higher capability of the given embedding methods. We train MARVEL on three datasets on half of slices with the other half slices as the unseen query slices. MARVEL achieves the best Top-K Hit Ratio across different values of K when compared to two baselines, demonstrating its potential to construct the ME reference atlas effectively.

### 3.4 Novel ME identification

In real-world applications, newly profiled samples may contain novel microenvironments (MEs) that are absent from the reference atlas. Consequently, a robust method should have the capability to identify these previously unseen MEs. However, this task has also not been addressed before. MARVEL is the first methods to address this task. Intuitively, those novel MEs should show minimal similar to any samples in the reference atlas. Therefore, we propose to the novelty score 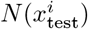 for each test sample 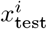 as

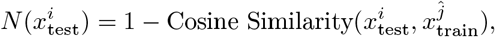

where 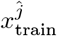 is the sample in the reference dataset most similar to 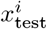. We define the ground-truth label 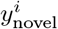 for each test sample as 1 if the sample belongs to a novel microenvironment (ME) and 0 otherwise.

To simulate this scenario, we separate the CODEX dataset based on the presence of the Follicle Tumor ME. Slices lacking the Follicle ME are designated as reference slices and form the training dataset *X*_train_, while slices containing Follicle ME are used as test slices, forming *X*_test_. The Follicle ME is annotated as the novel ME. We choose the Follicle ME because it is naturally absent in some slices due to the CRC subtype differences in the original dataset, making this novel ME biologically meaningful.

We evaluate the performance of our method using AUPRC and AUC-ROC to assess how well the novelty scores *N* can distinguish between novel and non-novel MEs. Compared with baseline methods, MARVEL achieves the best performance as shown in Fig 6a. These results first demonstrate the feasibility of our proposed new task (novel ME identification) in real biological applications. Additionally, they provide supporting evidence for MARVEL’s ME embedding, as MARVEL must both capture the information of seen ME categories and maintain generalization to unseen ME categories. This highlights MARVEL’s potential in complex real-world open set application scenarios.

**Fig. 4:**
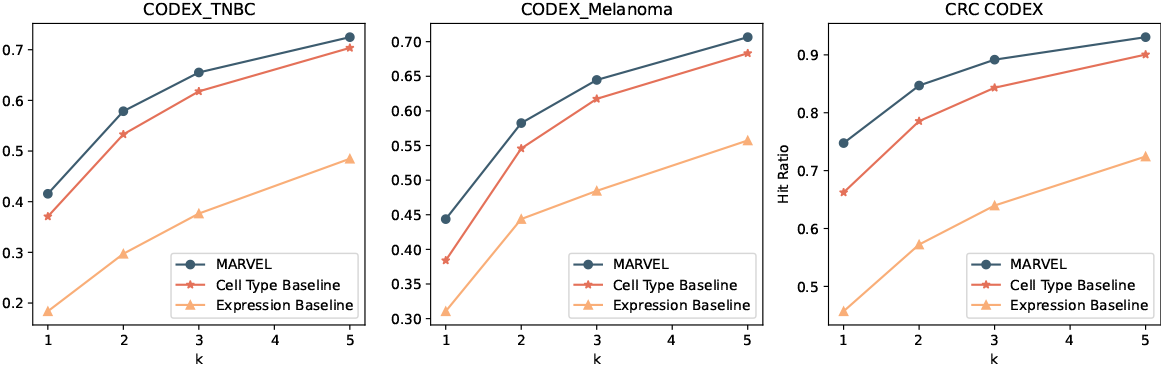
Inductive label query benchmarking results against various baselines across different values of K in Hit Ratio@K on three different datasets.

**Fig. 5:**
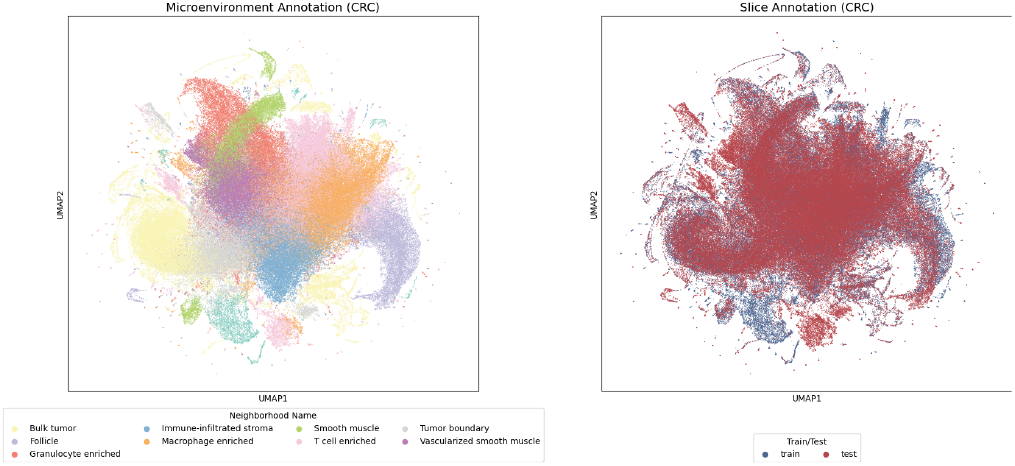
UMAP visualization of the learned representation with 0.1 labeled proportion for both reference and query microenvironments

**Fig. 6:**
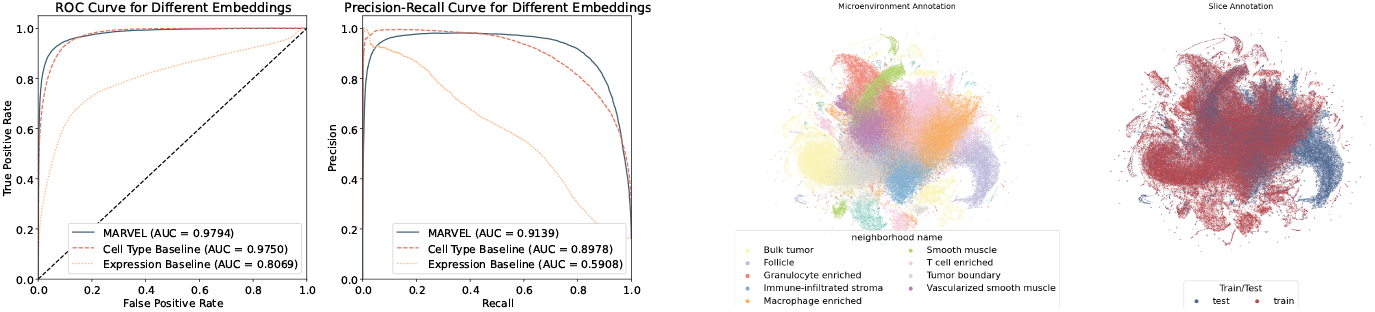
Novel microenvironment detection.Left, AUC-ROC and AUPRC benchmarking results against baselines for novel microenvironment identification in the CODEX CRC dataset. Right, Visualization of the learned ME representation for both query and reference datasets, where the query dataset contains a unique novel ME (Follicle) identified in the CODEX CRC dataset.

## 4 Conclusion and discussion

In the current spatial omics research domain, most works focus on unsupervised spatial domain clustering within individual slices to annotate microenvironments. However, these approaches often struggle to extend to multi-slice settings, limiting their ability to efficiently leverage existing labels for robust label transfer across large-scale datasets. Our proposed method, MARVEL, aided by the graph formalization of microenvironments and our proposed graph-level supervised contrastive learning, achieves efficient and robust construction of reference microenvironments with accurate annotations.

## 5 Code Availability

MARVEL source code is available at https://github.com/C0nc/MARVEL

## Notes

### Competing Interest Statement

The authors have declared no competing interest.

