## Supplementary material for "MARVEL: Microenvironment Annotation by Supervised Graph Contrastive Learning": Supp1

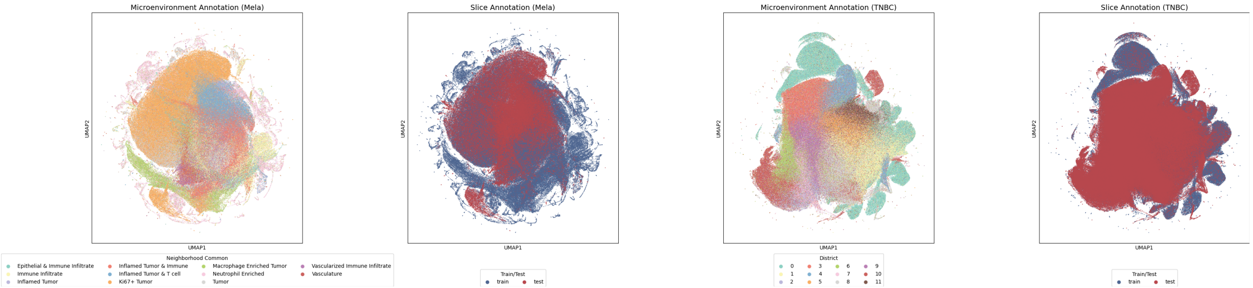


Figure 1: UMAP visualization of the learned representation with 0.1 labeled proportion for both reference and query microenvironments on remaining two datasets.

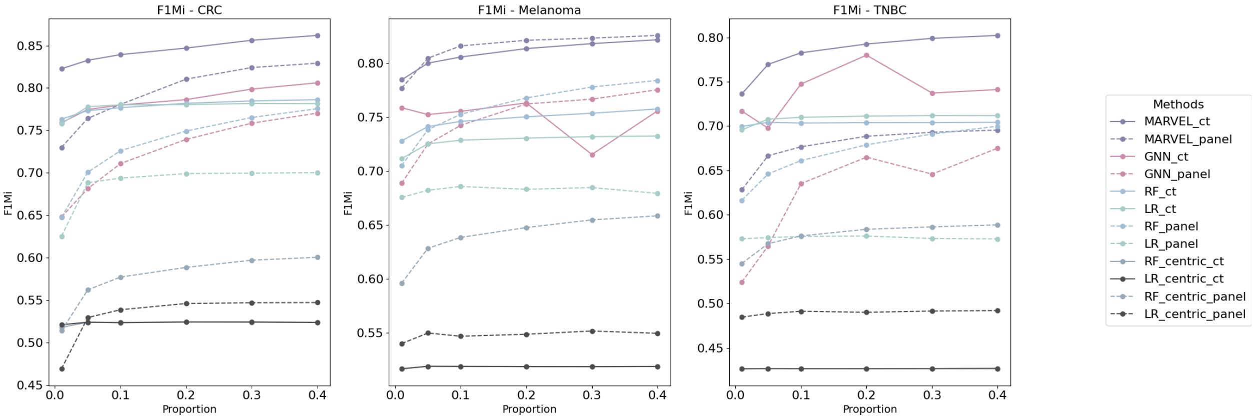


Figure 2: Transductive label transfer results (F1-Micro) across multiple baselines for different label proportions [0.01, 0.05, 0.1, 0.2, 0.3, 0.4] on three different datasets.
